# A probabilistic successor representation for context-dependent prediction

**DOI:** 10.1101/2022.06.03.494671

**Authors:** Jesse P. Geerts, Samuel J. Gershman, Neil Burgess, Kimberly L. Stachenfeld

## Abstract

The different strategies that animals use for predicting reward are often classified as model-based or model-free reinforcement learning (RL) algorithms. Model-based RL involves explicit simulation the future to make decisions while model-free strategies rely on learning associations between stimuli and predicted reward by trial and error. An alternative, intermediate strategy for RL is based on the “successor representation” (SR), an encoding of environmental states in terms of predicted future states. A recent theoretical proposal suggests that the hippocampus encodes the SR in order to facilitate prediction of future reward. However, this proposal does not take into account how learning should adapt under uncertainty and switches of context. Here, we introduce a theory of learning SRs using prediction errors which includes optimally balancing uncertainty in new observations versus existing knowledge. We then generalise that approach to a multi-context setting, allowing the model to learn and maintain multiple task-specific SRs and infer which one to use at any moment based on the accuracy of its predictions. Thus, the context used for predictions can be determined by both the contents of the states themselves and the distribution of transitions between them. This probabilistic SR model captures animal behaviour in tasks which require contextual memory and generalisation, and unifies previous SR theory with hippocampal-dependent contextual decision making.

## 1 Introduction

Humans and other animals are able to solve a wide variety of decision-making problems with remarkable flexibility. This flexibility is thought to derive from an internal model of the world, or ‘cognitive map’, used to predict the future and plan actions accordingly. Such model-based planning, which is flexible but slow, can be contrasted to model-free learning (Sutton & Barto, 1998). Model-free methods support rapid decision-making but are inflexible, as they forgo learning explicit knowledge about the environment’s dynamics in favour of learning simple associations between states or stimuli and cumulative long-term reward. An alternative, intermediate approach to RL takes the form of a representation of long-run state expectancies. This Successor Representation (SR; Dayan, 1993) combines aspects of model-based and model-free RL: planning with the SR can be done without explicit forward simulation and is therefore relatively computationally efficient. At the same time, the SR captures information about the environment structure that is independent of rewards, resulting in more flexible behaviour than simple model-free approaches when goals change (Russek et al., 2017).

The SR has recently gained considerable attention in psychology and neuroscience (Gershman, 2018; Momennejad, 2020). For example, a recent theoretical proposal suggests that hippocampal place cells, that fire when an animal visits a particular location in the environment (O’Keefe & Dostrovsky, 1971), in fact encode the SR (Stachenfeld et al., 2017). A different line of work has implicated dopamine neurons in the mid-brain, which are commonly thought to encode prediction errors about reward (Schultz et al., 1997), in computing prediction errors about state features, consistent with the prediction errors that are required for online learning of the SR (Gardner et al., 2018). These interpretations of the activity of hippocampal and dopaminergic cells suggest they are involved in a process of learning a representation of the environment that supports efficient prediction of future rewards.

Bayesian theories of learning suggest that animals not only learn to predict reward, but that they also estimate their uncertainty about these predictions. For example, previous work has demonstrated that a wide range of animal learning phenomena can be explained by probabilistic generalizations of simple model-free learning algorithms (Dayan & Kakade, 2001; Dayan & Yu, 2003; Gershman, 2015). These theories posit that, rather than learning a single (point) estimate for the weights used to approximate future reward, animals track a distribution over these weights or parameters. Such distributions contain additional information about the variance of the value function parameters, which reflect uncertainty, as well as information about the covariance between interdependent parameters. These uncertainty and interdependency terms can explain why animals learn more slowly in situations of low uncertainty (as evidenced by phenomena such as latent inhibition) and why they can learn about stimuli that are not currently present (as evidenced by phenomena such as backward blocking) (Dayan & Kakade, 2001; Gershman, 2015).

In addition to uncertainty about associations encountered within a particular context, animals might also represent uncertainty over which previously experienced task or context the current observations belong to. This type of uncertainty is of particular interest because the hippocampus is also implicated in context-dependent behaviour (Holland & Bouton, 1999). Previous theoretical work has suggested that the hippocampus performs this role in context-dependent behaviour by clustering observations as belonging to different latent causes (Fuhs & Touretzky, 2007; Gershman et al., 2010; Penny et al., 2013), with implications for memory updating (Gershman et al., 2014) and hippocampal remapping (Sanders et al., 2020).

In this paper, we introduce an extension to the SR model by augmenting it with an ability to track and use uncertainty. We first introduce a probabilistic interpretation of the SR that is learned using Kalman TD (Geist & Pietquin, 2010a), which will allow the appropriate balancing of prior beliefs and new sensory evidence to find an optimal learning rate. As in the value-learning case described by Gershman (2015), this allows for tracking uncertainty and covariance, explaining a set of animal learning phenomena that require learning about stimuli that are not currently present. We then generalise this approach to a multiple task or multiple context setting using a Bayesian nonparametric switching Kalman filter (Gershman et al., 2014), allowing the model to learn and maintain multiple task-specific SR maps and infer which one to use at any moment based on its sensory observations. In this definition, context reflects both the contents of sensory states and the transitions between them. This means that a new set of transition rules in the same environment can lead to a change of the context by which predictions are being made. We show that this probabilistic SR model accurately captures apparently contradictory animal behaviours in tasks which require contextual memory and generalisation.

## 2 Model description

This paper addresses the problem of how to deal with uncertainty when learning a predictive map. This predictive map takes the form of a Successor Representation (SR; Dayan, 1993), a representation of states in terms of the expected discounted future occupancy of each state, from the current state. The SR has been proposed to explain aspects of neural data in the rodent and human hippocampus (Brunec & Momennejad, 2022; de Cothi & Barry, 2020; Gershman, 2018; Momennejad, Otto et al., 2017; Russek et al., 2017; Russek et al., 2021; Stachenfeld et al., 2017). Our first contribution, which we initially described in Geerts et al. (2019), is to introduce a probabilistic SR, in which the agent’s belief about the parameters of the SR is expressed in terms of a distribution over possible SRs. This enables efficient learning by making use of the second-order statistics of predictions about future states or features, and can be used to understand a range of animal learning phenomena. Our second novel contribution is to extend the probabilistic SR to a probabilistic *hierarchical* SR in which the agent can switch between multiple SR maps when the environment, task, or context changes. This allows us to represent uncertainty within a context as well as over contexts. In this section we describe the pieces of the model in sequence. First, we describe how to handle uncertainty when learning the SR in a single environment using Kalman Temporal Differences (KTD) (Geist & Pietquin, 2010a; Gershman, 2015) applied to the SR. Next, we describe how to generalise this for multiple environments and context-dependent SR maps by using a nonparametric Switching Linear Dynamical System (SLDS) (Fox et al., 2011; Gershman et al., 2014; Murphy, 1998), that infers new maps when observations change drastically over time. Simulations using these models are presented in sections 3.1 and 3.2, respectively.

### 2.1 Background

We define an RL environment to be a Markov Decision Process consisting of *states s* the agent can occupy, *transition probabilities T_π_*(*s*′|*s*) of moving from state *s* to state *s*′ given the agent’s policy *π*(*a*|*s*) over actions *a*, and the reward available at each state, for which *R*(*s*) denotes the expectation. An RL agent is tasked with finding a policy that maximises its expected discounted total future reward, or *value*:

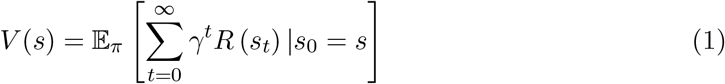

where *t* indexes time step and *γ*, where 0 ≤ *γ* < 1, is a discount factor that down-weights distal rewards. In classical model-free learning algorithms called temporal-difference (TD) learning (Sutton & Barto, 1998), *V* is learned directly through trial and error: each time a new state is encountered, *V* is updated proportionally to the difference between the expected and observed reward, the TD reward prediction error. However, such algorithms suffer from a lack of flexibility: when the reward function changes, model-free learners are slow to re-learn the new value function. Dayan (1993) proposed one solution to this problem, made possible by the fact that *V* decomposes into a dot product of the direct rewards *R* and a predictive representation *ψ*:

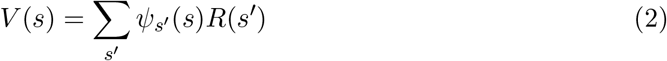

where *ψ*(*s*) is a vector with entries *ψ*_*s*_′/(*s*) containing the expected discounted future occupancy of state *s*′ along trajectories started in state *s* (see Figure 1A-B for a simple example):

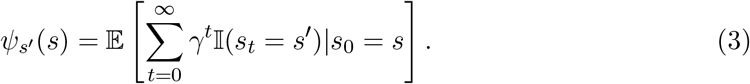

Factorising value into an SR term and a reward term permits greater flexibility because if one term changes, it can be relearned while the other remains intact (Barreto et al., 2016; Dayan, 1993; Gershman, 2018; Russek et al., 2017; Tomov et al., 2021). Since long term expectations about state occupancy can be slow to estimate, this lends particular robustness when reward is changing and transition dynamics are not.

**Figure 1:**
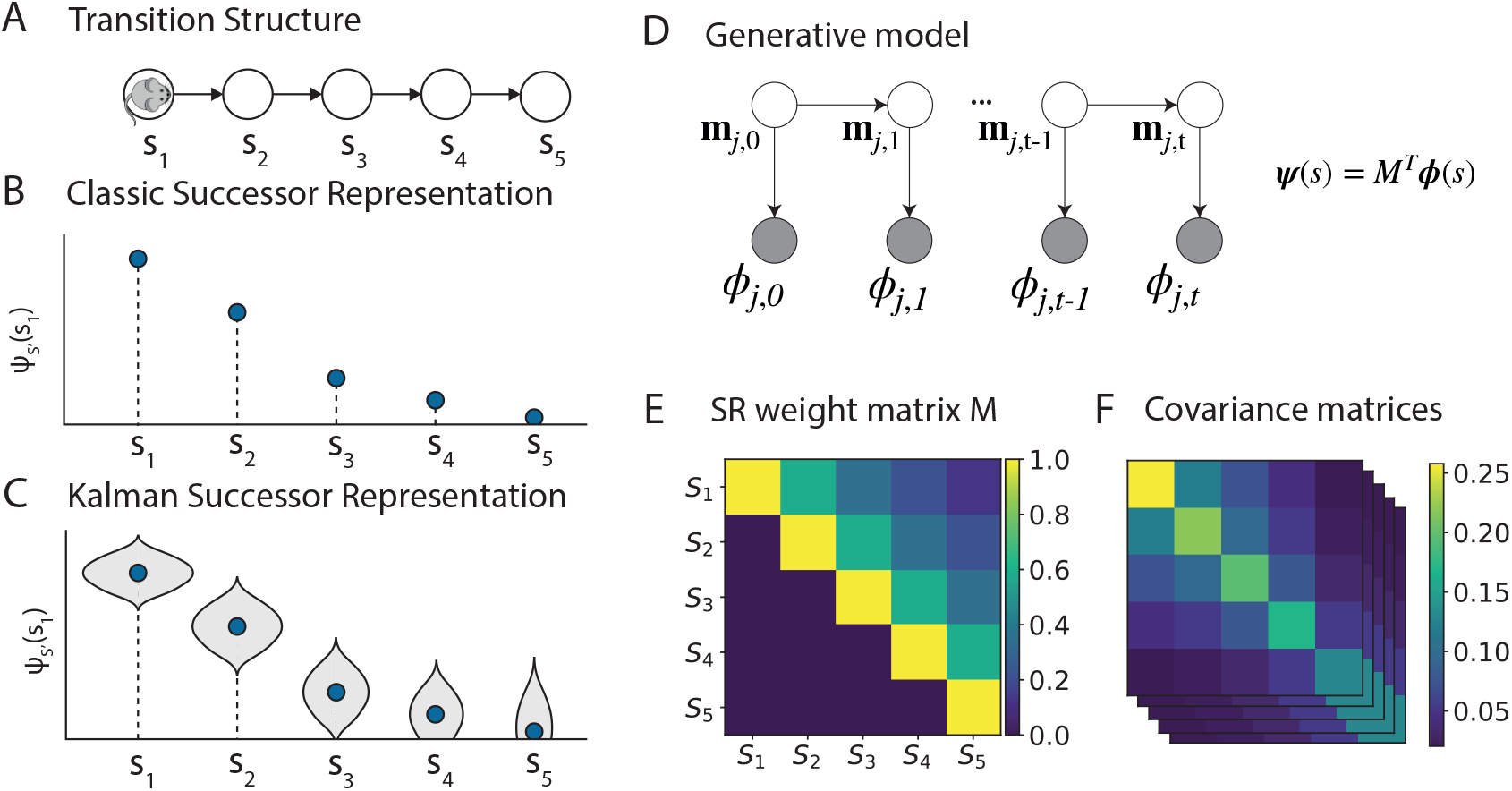
Model overview. **(A)** Transition structure of a sequence of five states. **(B)** Successor representation of state **s**_1_, corresponding to the expected discounted future occupancy given starting state **s**_1_ (blue dots). **(C)** In the Kalman SR model, a distribution over feature predictions is estimated: in addition to the mean (blue dots), the variance of each successor feature is estimated (grey shaded region shows distribution). Note that the distribution includes estimates below zero, outside the limits of this figure. **(D)** The Kalman SR generative model’s graphical structure. *ϕ_j_*, *t* denotes the *j^th^* feature of state *s_t_*, **m**_*j*_,*t* denotes the *j^th^* column of the (latent) SR weight matrix shown in (E), at time **t**. *ψ* (**s**) denotes the vector successor representation at state **s**. **(E)** The SR weight matrix corresponding to the transition structure in (A). The relation between the weight matrix *M* and the current representation *ψ* is given in the inset equation in (D). **(F)** In addition to a mean estimate of *M*, Kalman SR represents the uncertainty over SR weights with a set of covariance matrices corresponding to the columns of *M*.

The SR can be generalized to continuous states by representing states using a set of feature functions *ϕ_j_* (*s*). In this case, the SR is referred to as Successor Features (SFs, Barreto et al., 2016), and encodes the expected feature values:

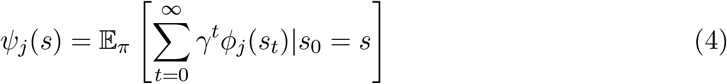

Specifically, *ψ_j_*(*s*) denotes the expected discounted future occurence strength of feature *j* from state *s*. In this linear function approximation case, the reward is given by the dot product

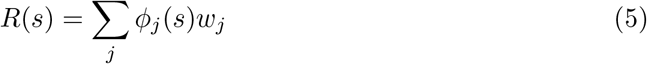

where *ϕ*(*s*) are the state features and **w** are weights parameterising the reward function.

The decomposition of value (Equation 2) is then re-written as:

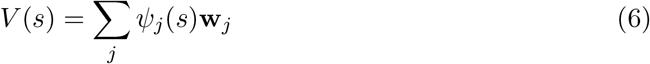

In the special case where the state space is finite and *ϕ* is a tabular representation of the state (i.e. Figure 1A, where states are discrete and represented as one-hot vectors), Equations 4 and 6 reduce to Equations 3 and 2. The contents of the feature vector can be arbitrary, and will in this paper depend on the particular task being modelled. We model this feature-based SR as 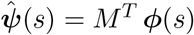, where *M* is a weight matrix in which each entry *M_ij_* indicates the extent to which feature *i* predicts feature *j* (Figure 1E). We thus assume for now that the agent has access to a state representation *ϕ* such that a linear mapping exists from features of each state to the discounted future occurrence of those features, given the start state. Seen as a single layer of a biological neural network, each column of *M* comprises a vector of input weights of one SR-encoding neuron *ψ_j_*, and the vector *ψ*(*s*) gives the population activity of SR-encoding neurons. To avoid cluttered notation, we will denote a column by **m**_*j*_ = *M*_:,*j*_ so that

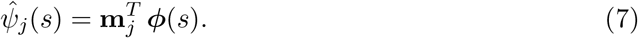

The factorised representation of the value function in equation 6 means that two quantities have to be learned: the reward weights **w** and the SR *ψ* (note that, although we do not model this here, *ϕ* can be learned too; see e.g. Hansen et al., 2019). We track the reward weights with a Kalman Filter (Gershman, 2015). The SR *ψ* can be learned with TD learning, in which the SR is updated according to a TD state prediction error reflecting the difference in estimates of *ψ*(*s*) and estimates of 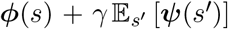 (Dayan, 1993; Gardner et al., 2018). TD learning of the value function (using reward rather than state prediction errors) is a popular model of learning in the striatum (Schultz et al., 1997), and TD algorithms can be implemented in biologically realistic spiking networks (Bono et al., 2021; Brea et al., 2016; Frémaux et al., 2013).

### 2.2 Probabilistic Successor Features

In its original formulation, the SR computes a point estimate of *ψ_j_* from the values of the SR weight matrix *M*. Our first step is to replace this with a probabilistic description of the SR. This probabilistic SR explicitly represents uncertainty. We model each column **m**_*j*_ of the SR weight matrix *M* as a set of random variables. Under this interpretation, the animal implicitly assumes there is a true, hidden set of SR parameters **m***j*, which predict each new noisy observation via a generative model. The animal’s goal is to invert this generative model in order to infer a distribution over the SR weights from observations. More precisely, from a sequence of observations *ϕ*_1:*t*_, the agent can infer information about the hidden SR weights using Bayes’ rule:

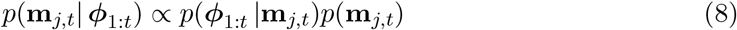

This idea, in the form of a Kalman Filter (Kalman, 1960), has previously been applied to learning value functions (Geist & Pietquin, 2010a; Gershman, 2015) and readily applies to the SR. In our SR model, each column **m**_*j*_ of *M* is modelled as a vector-valued random variable, with dimensionality *N_ϕ_*.

#### Generative model

Performing inference requires specifying a probabilistic generative model relating the hidden SR weights to the animal’s observations (Figure 1D). It consists of a *prior* on each column of the SR matrix **m**_*j*,0_, an *evolution* equation describing how these hidden SR vectors evolve over time and an *observation* equation describing how the hidden SR relates to observations. The observation equation follows directly from the Bellman equation, with additive Gaussian observation noise 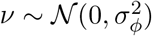:

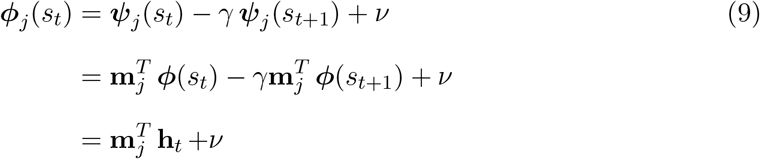

where we have defined **h**_t_ = *ϕ*(*s_t_*) – *γϕ*(*s*_*t*+1_) to be the discounted temporal difference in feature observations. We assume, in other words, that each successor feature *ψ*_j_(*s_t_*) is a noisy linear function of the current features (see Appendix B for additional analysis). For the evolution equation, our generative model follows a Gaussian random walk allowing the weights to change incrementally over time. We also assume a Gaussian prior on the weights. Together, these form the following probabilistic generative model (shown in Figure 1D):

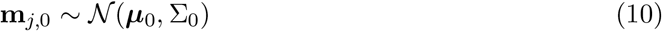

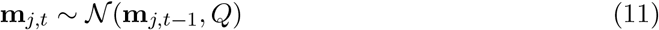

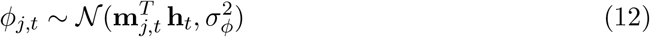

where ***μ***_0_ is the prior mean, ∑_0_ is the prior covariance matrix, 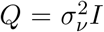 is the (diagonal) transition noise covariance matrix with transition noise variance 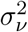 and 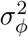 is the observation noise variance. Parameter values used in the simulations are given in Table 1.

**Table 1:** Parameter settings used in the simulations.

| Symbol | Value |
| --- | --- |
| $\alpha$ | 1.0 |
| $\beta$ | 1.5 |
| $\gamma$ | 0.9 |
| $\sigma_\phi^2$ | 1.0 |
| $\sigma_\nu^2$ | 0.001 |

#### Inference

Our goal is to estimate the parameters **m**_*j*_ such that they satisfy 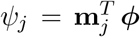, for each successor feature *j* given the observation *ϕ*. Since the generative model described above is a linear-Gaussian dynamical system (LDS), we can perform exact inference on these SR weights by combining the Kalman filter equations with TD learning. Estimating a distribution over SR weights involves adjusting the mean estimate **m**_*j*_ using a temporal difference learning rule, but now taking into account the relative covariances of the weights, ∑, via the Kalman gain ***κ***, an adaptive, feature-specific learning rate (Figure 1E-F). This allows for a closed-form update of a *posterior distribution* over the weights (Figure 1C):

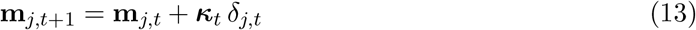

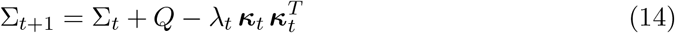

where 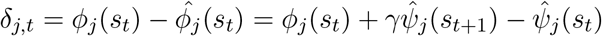 is the successor prediction error for feature *j*, 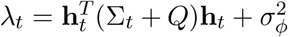 is the residual variance, and ***κ**_t_* is the Kalman gain is given by:

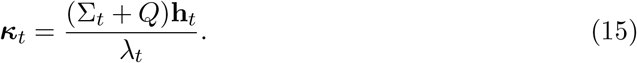

Importantly, this learning rate is feature-specific and dependent on the covariance.

The Kalman Filter’s covariance-dependent learning rate gives rise to several learning phenomena that have been previously explored in the literature. When the uncertainty about the hidden weights **m**_*j*_ is low compared to the uncertainty of the observation (low variance, in the diagonals of Σ), the posterior should be close to the prior, resulting in a lower learning rate. Under high uncertainty (high variance), newly incoming observations should be weighted as more informative and the learning rate should be high. When there is non-zero covariance between a set of weights, these weights are updated together because they share the same ***κ**_t_*. This permits non-local updating of parameters; that is, parameters for features not present in the current observation may be updated if these parameters have a known covariance with parameters in the current observation. In standard TD learning, the update equation would have the prediction error multiplied by the activity of the feature neuron and a scalar learning rate *η*

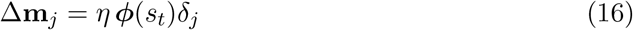

Replacing the first two terms with ***κ*** means that learning can occur without feature neuron activity (c.f. equation 13).

### 2.3 Inferring SR and context simultaneously

By design, Kalman filter models evolve smoothly, and do not capture situations where the hidden variable undergoes large sudden changes or jumps (Figure 2). However, such sudden changes in the environment might occur when the animal switches to a different context, or returns to an old one.

**Figure 2:**
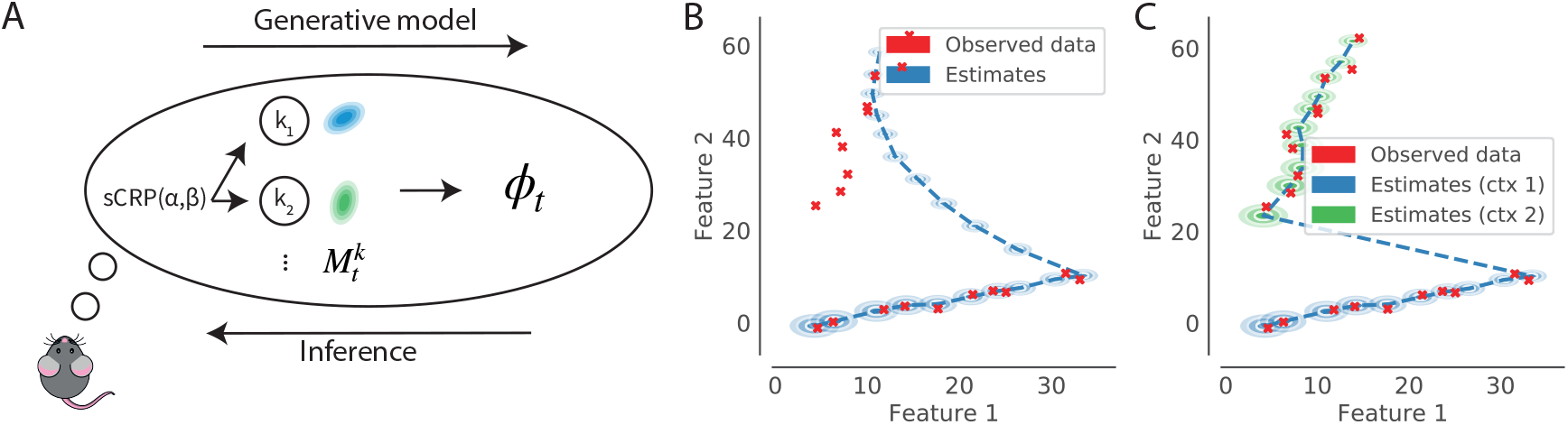
Switching Kalman filter model illustration. **(A)** In the infinite switching Kalman filter generative model, a context *k* is drawn from a sticky Chinese Restaurant Process (sCRP) prior. The currently active context selects one of infinitely many possible linear-Gaussian models to pass through to the observations, *ϕ_t_*. Given this generative model, the animal’s goal is to infer both the SR parameters, *M* and the discrete context variable, *k*. **(B)** A single Kalman filter does not account for large jumps in the hidden variable that is tracked (ellipses show the posterior distribution at each time step). **(C)** A switching Kalman filter deals with large prediction errors by assigning them to a new context (posterior distributions are colour-coded with the inferred context). (B) and (C) show a series of observations of a pair of features over time (red crosses, feature 2 increases monotonically with time) and their estimated posterior distribution (shaded ellipses) under context 1 in (B) and with a model that can switch to a new context 2 when prediction errors are large.

We can account for these jumps with a “switching LDS”, which posit that there is a collection of different modes or contexts, in which each context is associated with its own LDS. This means that our model switches between different SR maps *M*^*k*^ that correspond to different contexts *k*. Since there are infinitely many possible contexts, we use a nonparametric switching LDS (Fox et al., 2011) which allows the number of inferred contexts to grow as more observations are made. This generative model corresponds to that used in Gershman et al. (2014) to model memory updating, with the difference that here the continuous hidden state will be the SR. Thus, an SR-context is chosen if it correctly predicts observations. If there are no such predictive contexts, a new SR is created.

#### Generative model

In the generative process this model assumes, a context *z_t_* is first drawn from a sticky Chinese restaurant process (sCRP) prior. A CRP prior allows for a potentially infinite number of contexts, but tends toward fewer contexts by proportionately assigning observations to contexts that already explain more observations. The “sticky” CRP has an additional bias to remain in the current context. The CRP prior is written as:

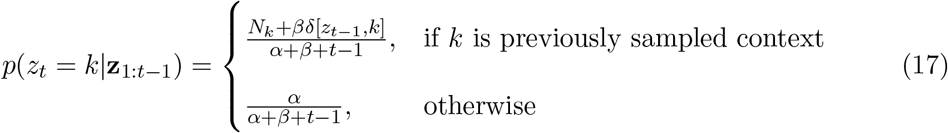

where *N_k_* is the number of observations previously assigned to context *k* and *δ*[*x,y*] = 1 if *x* = *y* and 0 otherwise. The concentration parameter *α* controls the propensity to create new modes and the “stickiness” parameter *β* determines how likely the model will stay with the current context.

After choosing a context, the generative model proceeds by evolving the state variable for each previously active context *k* according to the evolution equation of the LDS: 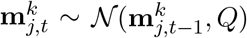. If *z_t_* is a new context, a new SR is first drawn with columns 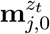 drawn from a Gaussian prior: 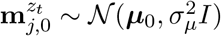. Finally, a sensory observation *ϕ_t_* is emitted from the currently active context *z_t_* using the observation equation: 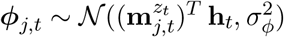.

#### Inference

When there is uncertainty about the context, inference requires marginalising over all possible context histories **z**_1:*t*_:

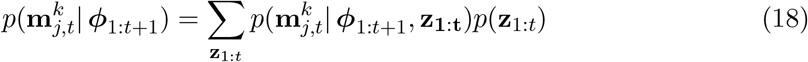

If there are *K* modes, the posterior at time *t* will be a mixture of *K^t^* Gaussians, one for every possible history *z*_1_,… *z_t_*. Exact inference is intractable under this exponentially growing number of modes. We therefore use the particle filter (Fearnhead & Clifford, 2003; Murphy, 1998), which approximates the posterior distribution over contexts at each time step using a set of weighted particles that are updated sequentially as new observations are obtained.

The particle filter samples a context assignment *z_t_* for each particle {*l*} from the sCRP prior (equation 19), and to use the Kalman filter equations to estimate each 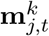, with Kalman gain:

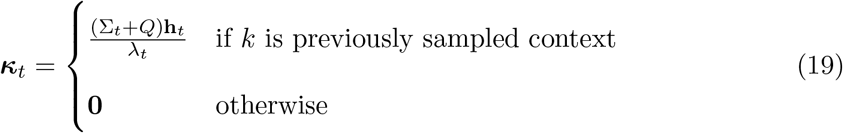

Each particle will represent a possible history of context assignments, and the posterior over contexts is obtained by averaging. The particles’ weights are updated according to 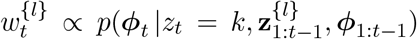, which corresponds to the one-step ahead predictive density, or the likelihood of the next observation, given by:

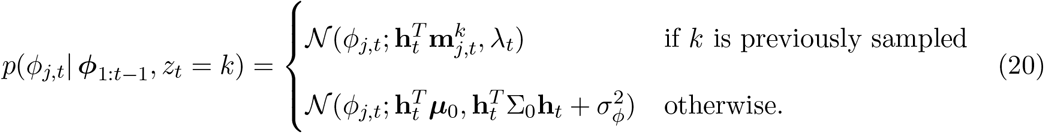

This corresponds to choosing contexts according to how well they predict observations: each previously used context *k* makes a prediction centred on 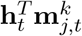. Thus, the log-likelihood of any existing context will be inversely proportional to the magnitude of the SR prediction error for that context, || *ϕ_t_*–*M^T^***h**||^2^, meaning that very large prediction errors will likely lead to the inference of a new context. Furthermore, since the variance of a context grows with the amount of time since its last occurrence, older modes will be more tolerant of prediction errors. The intuitive explanation for this is that if the animal has not seen a context for a long time, its certainty about the details of the events will have deteriorated.

In summary, the Kalman SR algorithm operates by predicting the occurrence of a feature using the weights 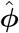 and updating its prediction based from observation. Updates are sensitive to the covariance between predicted features, such that the magnitude of the update is informed by uncertainty and covarying features are simultaneously updated (permitting learning about features not currently experienced). We model context-dependent SR learning using a nonparametric switching Kalman Filter, in which the SR diffuses gradually until it jumps to either a previously activated context, or to a new one. This allows us to model how uncertainty over the active context modulates inference using the SR. The Bayesian view of the SR outlined here allows us to reconcile and reinterpret some results in the animal learning literature, which we will describe in the following section.

## 3 Results

In the first part of this section, we will discuss results that follow from the single Kalman filter SR described in Section 2.2. In the second part, we will discuss experimental predictions relating to the switching context model described in Section 2.3.

### 3.1 Kalman SR simulations

#### Facilitation and latent inhibition in contextual fear conditioning

Prior work on contextual fear conditioning in rodents looks at the conditioned response (freezing) following an aversive stimulus (a small foot shock) received in some new environment. It has been shown that the a stronger conditioned response is evoked if animals are able to explore the new environment for several minutes before the first shock, a finding known as the ‘context pre-exposure facilitation effect’ (Fanselow, 2010). As pointed out by Stachenfeld et al. (2017), a predictive model such as the SR can account for this: during pre-exposure, the animal explores and learns a predictive representation of the context such that subsequent value learning is rapidly propagated across the environment. However, context pre-exposure facilitation stands in apparent contrast to “latent inhibition”, which refers to the finding that pre-exposure to a conditioned stimulus (CS) typically *impairs* the acquisition of a conditioned response. This latent inhibition effect has been taken as evidence for the assertion that animals are Bayesian learners: as the pre-exposed cue is presented repeatedly, the animal’s uncertainty about the expected reward associated with that cue decreases, resulting in slower subsequent learning (Gershman, 2015).

Latent inhibition and facilitation are thus opposing effects that could both be driven by context pre-exposure. Kiernan and Westbrook (1993) showed that there is an intriguing time course to these phenomena: a brief amount of pre-exposure facilitates subsequent learning, whereas extended pre-exposure to the environment inhibits further associative learning (Figure 3A). The Kalman SR model naturally captures this non-monotonic relationship, as both facilitation (driven by the SR) and inhibition (driven by the Kalman filter) occur during pre-exposure. As the animal explores the environment, there should be facilitation early on as the SR is learned. However, this should be followed by inhibition after extensive training because reduced variance in the estimates of the reward weights **w** results in a decrease in Kalman gain, as shown in Figure 3B.

**Figure 3:**
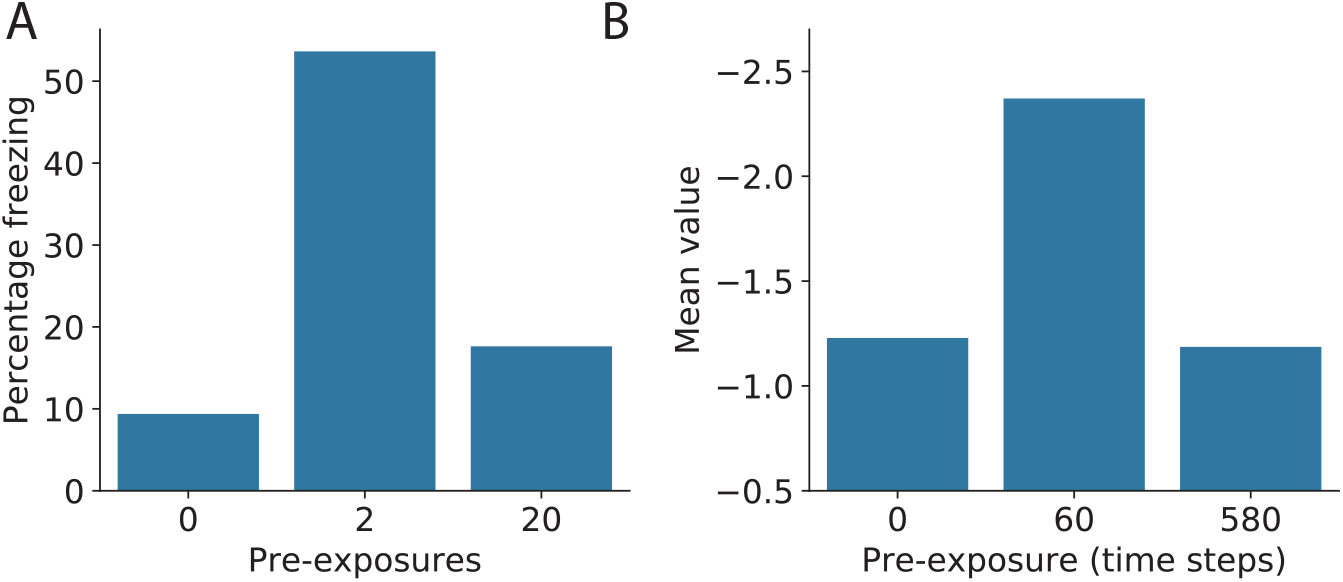
A brief amount of pre-exposure to an environment facilitates subsequent learning but extended pre-exposure impairs learning. **(A)** Behavioural data from Kiernan and Westbrook (1993). Mean percentage freezing scores in the shocked environment E1 for the groups receiving no, brief, or extended pre-exposure to E1 and exposed to a T1-T2 interval of either 7 sec or 60 sec. **(B)** Under the Kalman SR model interpretation, exploring the environment during pre-exposure allows a predictive representation to be learned. Since value is computed by multiplying the SR by the reward function, this means that longer pre-exposure initially facilitates learning the negative value in the environment. Prolonged pre-exposure, however, causes a decrease in uncertainty and therefore in Kalman gain, inhibiting further learning. Simulation results show the mean value estimated by the model.

#### Transition revaluation

A key prediction of standard temporal difference SR learning is that “reward revaluation” (changes in the reward at each state) should be easier to acquire than “transition revaluation” (changes in the transition probabilities between states), since the latter requires propagating state occupancy predictions to distal states. This is because temporal difference learning updates the SR only for experienced states, even though predictions from previous states are also ultimately affected by the change. Further experience is needed to update all affected states (note that eligibility traces could address this issue, but only for situations where the affected states are experienced in the same episode). Momennejad, Russek et al. (2017) tested whether or not this is the case in human learning. In their experiment, participants learned about two sequences of states leading up to a reward (see Figure 4A). In the next phase, half of the participants were exposed to a transition revaluation condition, observing novel transitions leading up to the reward. The other half experienced “reward revaluation” in the form of novel reward amounts at the final states. Importantly, the novel experiences in Phase 2 did not include starting states 1 or 2, meaning that under a classical SR model, the SR for states 1 and 2 would not be updated.

**Figure 4:**
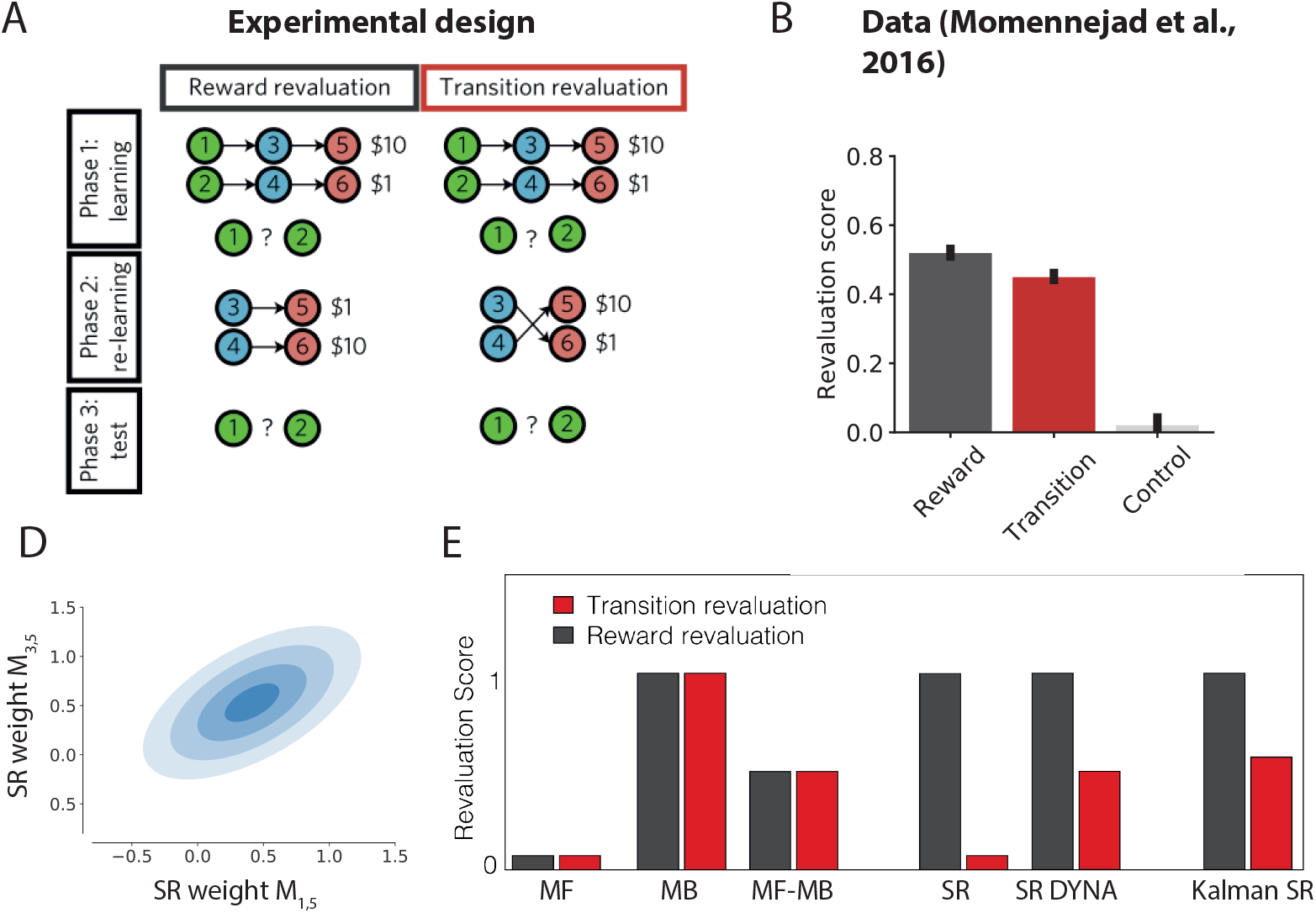
Revaluation experiment of Momennejad, Russek et al. (2017). (A) Experimental design. In an initial learning phase, participants learned sequences of states, associated with high ($10) or low ($1) rewards. During a second re-learning phase, either the rewards associated to the two terminal (red) states were swapped (reward revaluation) or the transitions from the middle (blue) to the terminal states were swapped (transition revaluation). (B) Human participants’ revaluation scores (Momennejad, Russek et al., 2017). (D) The joint distribution over weights *M*_1,5_ and *M*_3,5_ shows a positive covariance induced by the first phase of learning, which explains the revaluation from state 1 to 5. (E) Predicted revaluation scores (change in rating (*V*(1) – *V*(2)) between phase 1 and 3 for different algorithms.

While participants were significantly better at reward revaluation than transition revaluation, they were capable of some transition revaluation as well (Figure 4B). Accordingly, the authors proposed a hybrid SR model: an SR-TD agent that is also endowed with capacity for replaying experienced transitions (Figure 4E). This permits updating of the SR vectors of states 1 and 2 through simulated experience. Note that this pattern of results cannot be explained by a simple model-free (MF) or model-based (MB) strategy, or by a simple hybrid (MF-MB), as MF methods are equally unable to do either form of revaluation and MB methods equally able to do both (Figure 4E).

Simulating this experiment with Kalman SR shows that the model can account for the partial transition revaluation without explicit simulated experience. Kalman SR correctly learns the SR matrix after phase 1 as well as an estimate of the covariance between features, Σ. Unlike standard temporal difference methods, Kalman TD uses the covariance matrix to estimate the Kalman gain and uses that to update the SR non-locally. This means that after seeing 3 → 6, it updates not just *ψ* (3) but also *ψ*(1) because these entries have historically covaried (and similarly for *ψ*(4) and *ψ*(2)).

#### Reward devaluation of preconditioned cues

Our model similarly captures the findings of Hart et al. (2020), which show that responding to preconditioned cues is sensitive to devaluation, a hallmark of model-based learning. The particular experiment of interest here started with a preconditioning phase, during which associations were learned between pairs of neutral (i.e. nonrewarding) stimuli, followed by a conditioning phase during which a neutral stimulus was paired with a food reward (Figure 5A). After this conditioning phase, the food reward was devalued in one group but not another, by pairing it with sickness-inducing lithium chloride (LiCl). A key finding of this experiment was that healthy animals’ responding to the unblocked preconditioned cues (C) was sensitive to the subsequent devaluation of the food reward (Figure 5B, see also Hart et al., 2020). Thus, reward devaluation can alter stimulus-stimulus associations that were learned through the activation of dopamine neurons. Furthermore, the value inference depended on an intact orbitofrontal cortex (OFC) during the preconditioning phase (see Discussion).

**Figure 5:**
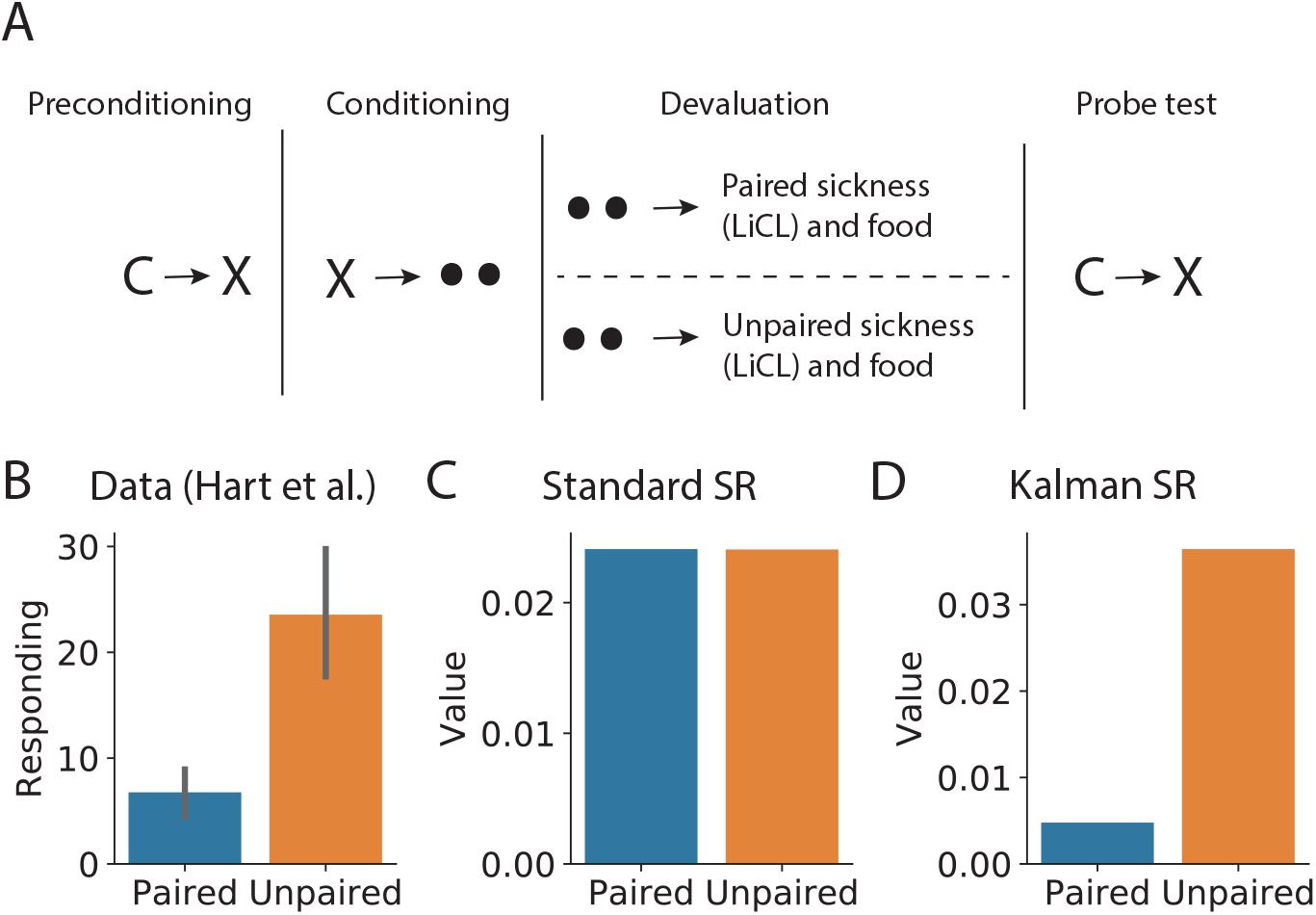
**(A)** Experimental design (Hart et al., 2020). During the initial preconditioning phase, one neutral stimulus always precedes a second (*C* → *X*). During the conditioning phase, the second stimulus is paired with a food reward. After the conditioning phase, the food is paired with lithium chloride (LiCl) to induce sickness in one group of animals. Letters denote different neutral stimuli, black circles indicate food reward. **(B-D)** Data and simulation results show that, like animals and unlike TD-SR, Kalman SR shows sensitivity to devaluation in this paradigm. Data in (B) replotted from Hart et al. (2020), error bars show SEM.

The SR naturally accommodates many preconditioning phenomena, because the separate representations of stimulus-food predictions and their valence allows for flexible revaluation (Gardner et al., 2018). However, in the experiment shown in (Figure 5A), the food reward was paired with illness in the absence of any of the neutral stimuli introduced in the preconditioning stage. This means that a standard SR agent would not be sensitive to the reward devaluation (Figure 5C). This is because in the TD SR, only stimuli that directly predict reward will change value after devaluation, and C was never directly associated with food. This was also observed by Gardner et al. (2018), who simulated a very similar task with an SR model endowed with the ability to simulate offline experience (see Appendix A, Figure A1). This latter model showed the same sensitivity to devaluation that was shown by the animals.

Like the animals in Hart et al. (2020), Kalman SR was sensitive to the reward devaluation paradigm. During the pre-conditioning phase, a positive covariance between C and X is learned, which means that during conditioning, C becomes directly associated to the food (equation 13). Subsequent devaluation thus directly affects C as well as X. This permits long range temporal credit assignment without explicitly necessitating hand engineered features or simulated sequential experience.

### 3.2 Switching context model simulations

#### Contextual memory

The context-switching model allows us to explain an intriguing finding in contextual fear-conditioning (Figure 6). Winocur et al. (2009) exposed rodents to a CS-US pair in context A, and subsequently tested in either context A or B, after either short (24 hr) or long (28 day) delays (Figure 6A). They found that the animals learned the association in context A, but did not generalize their conditioned responding to context B after the short delay (Figure 6D). However, the level of generalization increased with the delay interval.

**Figure 6:**
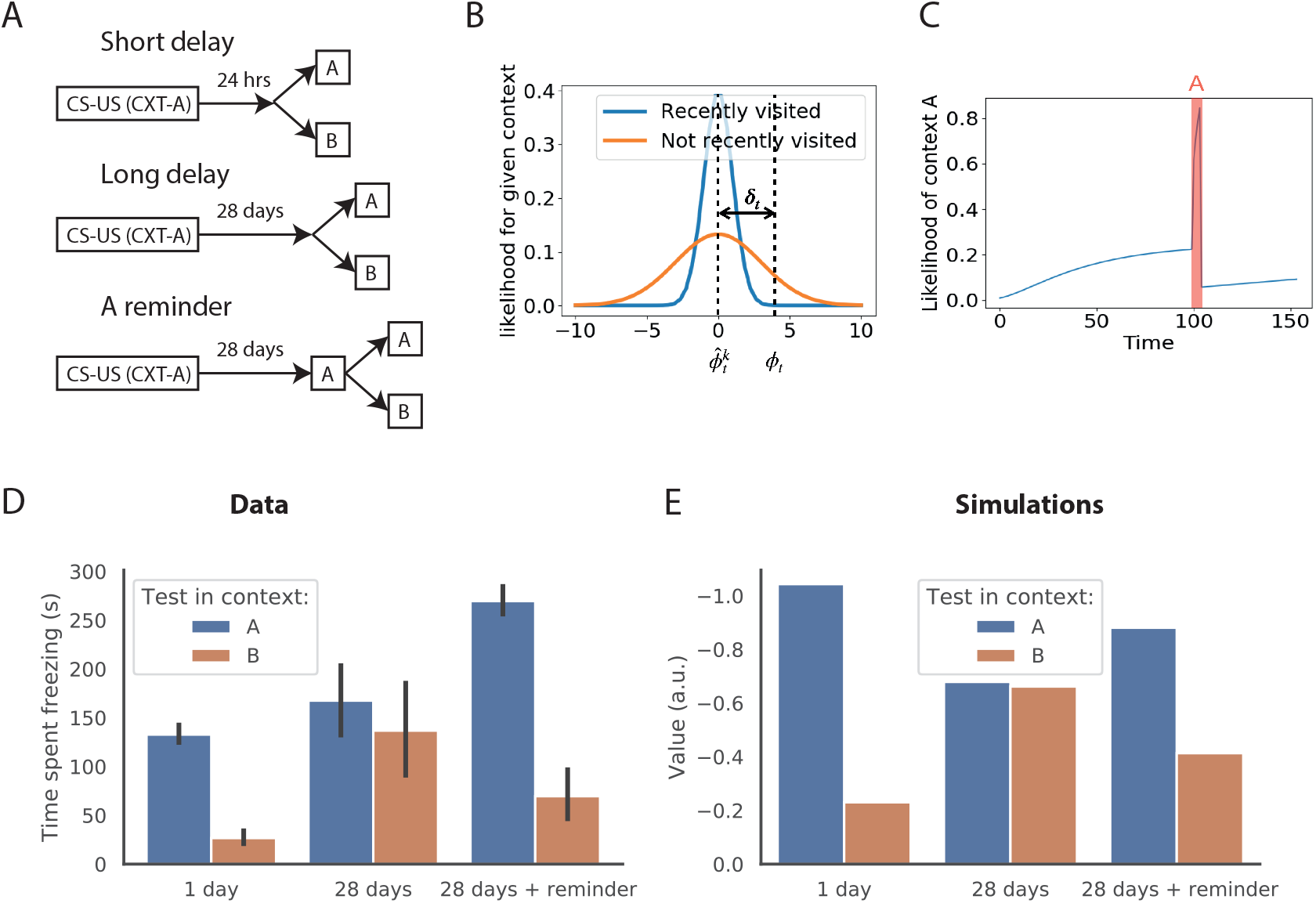
Contextual memory experiment by Winocur et al. (2009). **(A)** Experimental design. In the short delay condition, animals were conditioned in context A, and then tested in context A and a different context B, 24 hours later. In the long delay condition, there was a 28 day delay between conditioning and testing. In the reminder condition, animals were briefly reintroduced to context A, without administering the CS or US, before testing. **(B)** In the model, each context’s likelihood is a Gaussian centered on that context’s predicted observation *ϕ*. The larger the SR prediction error for that mode, the lower the likelihood. Sub-models for contexts that have not been active for a long time will have higher variance around the predicted mean and be more tolerant to prediction errors. **(C)** A reintroduction to the original context (red shaded region) reduces that context’s model’s variance, and hence it reduces the likelihood of inferring the context given a large prediction error. **(D)** Data replotted from Winocur et al. (2009) showing the time spent freezing in response to the CS in different conditions. **(E)** Simulation results showing the state value estimate when the CS is shown in different conditions. As in the data, value is increasingly generalised to context B, but a reminder of the original context restores context specificity.

Furthermore, when animals were briefly exposed to the training context, as a reminder prior to test in the second context after the long delay, the generalization decreased again (Figure 6D). Thus, cross-context specificity decreases with time but can be restored with a reminder of the context.

Our switching Kalman SR model explains this result in terms of switching between contextual representations. When the distance between the observation and the predicted observation given a context (successor prediction error: 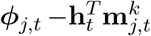) is large, the model will assign a low likelihood to that context. If the posterior probability of every currently active context is low, the model will be likely to assign the observation to a new cluster, initiating the use of a new, separate predictive map. Furthermore, since the variance of clusters that have not been visited for a while keeps growing (as can be seen in the Kalman filter updates above), old clusters will be more ‘tolerant’ to prediction errors, i.e. their likelihood will be larger, even for larger distances (Figure 6B). A re-exposure to the original context will reduce the variance again, restoring the sensitivity to prediction errors (Figure 6C). Thus, the model predicts that learning is highly context-specific early on but will lose context-specificity with time because of growing within-context uncertainty. Furthermore, because the Kalman filter’s covariance updates do not depend on the reward outcomes, mere re-exposure to the features of context A should restore the context-specificity of the learned predictions, thus recapitulating the results observed by Winocur et al. (2009) (Figure 6E).

#### Contextual generalization

In addition to elapsed time, the amount of contextual generalization is also dependent on the amount of initial exposure to the original context. This was shown by Kiernan and Westbrook (1993) in a follow-up experiment similar to that described in the previous section (Figure 7). The amount of context pre-exposure was again varied, and the propensity of animals to show conditioned responding was now recorded both in the original environmental context and in a novel environmental context, to test whether animals would generalize their responding in the original environment to a novel context. Recall from the previous section that these authors showed a non-monotonic effect of pre-exposure duration *within* a context, whereby pre-exposure to a context first facilitates, then inhibits learning (Figure 3). In contrast, increasing pre-exposure duration monotonically *decreases* the amount of generalization of the fear response to a second context (c.f. black and white bars Figure 7A). Under our model, this can be explained because increased exposure to the context results in a sharper posterior over the SR and reward weight parameters (Figure 7B). This *reduces* contextual generalization because the likelihood of the original context 1 will be low in context 2: since context 1’s SR is represented with a high precision, small differences in context 2 will not cause it to be grouped with context 1.

**Figure 7:**
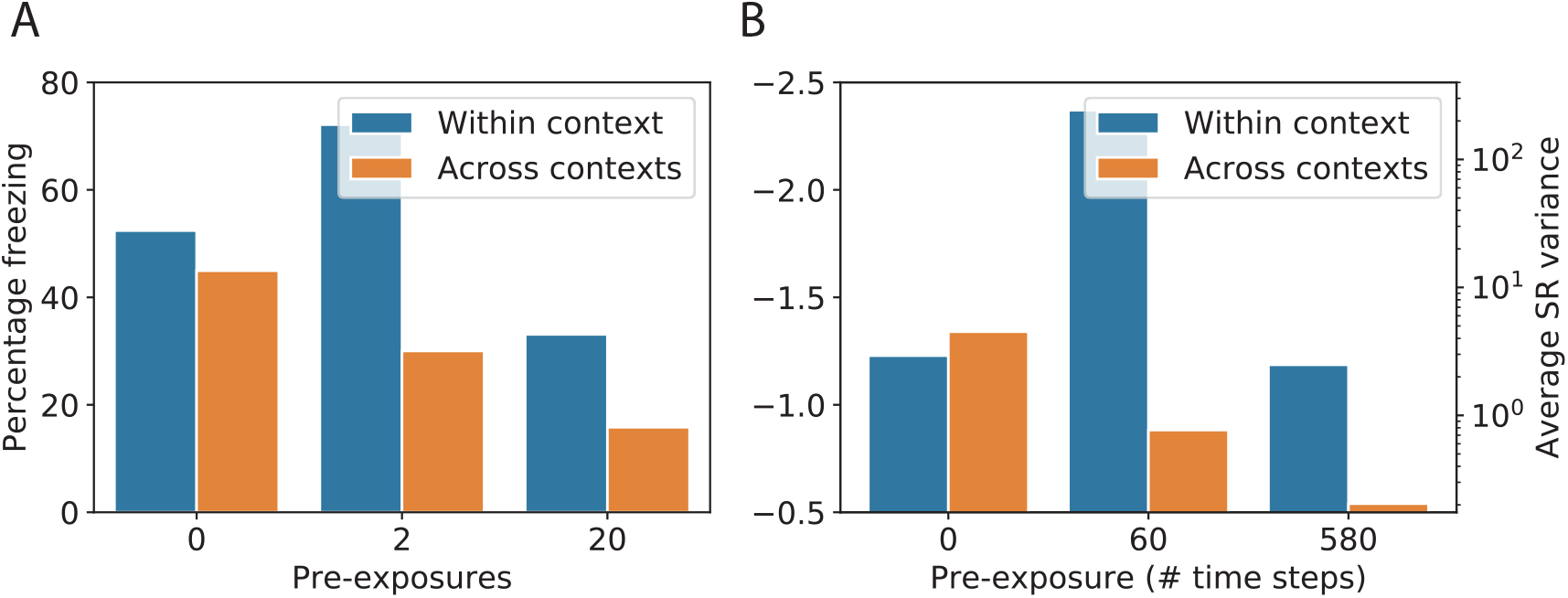
**(A)** Contextual discrimination data from Kiernan and Westbrook (1993, left). Black bars show conditioned freezing responses after conditioning for animals pre-exposed 0, 2 and 20 times to environment 1. White striped bars show the conditioned response to the same cue in a different environmental context. The same effect is shown for two experimental conditions in which the T1-T2 interval (i.e. the time between entering the environment and the presentation of the CS) was varied. **(B)** Model simulation results showing the negative value estimated by the model after conditioning as a function of pre-exposure time in black. In white, the average variance of the Kalman SR model is shown.

## 4 Discussion

The SR constitutes an efficient, flexible middle ground between model-based and model-free RL algorithms by separating reward representations from cached long-run state predictions. Here we introduce a probabilistic SR model using Kalman temporal differences that supports principled handling of uncertainty about state feature predictions and interdependencies between these predictions. This model is extended to a switching Kalman filter that switches between different modes or contexts. We show that these models capture human and animal behaviour in settings of context pre-exposure, transition revaluation and contextual generalisation and memory.

### 4.1 Potential roles for replay

An attractive feature of models such as the Kalman filter that track the covariance between different weights is that this allows for retrospective revaluation. This feature has previously been used to explain learning phenomena such as backward blocking (Dayan & Yu, 2003; Gershman, 2015) and more sophisticated Kalman filter models in which stochasticity and volatility parameters are estimated from data can further extend explanations to effect like the robustness of partial reinforcement (Piray & Daw, 2021a). Applied to SR learning, we have shown that this can extend to re-evaluating states after a change in the transition structure (Figures 4, 5 and A1). These effects have been explained in the past by positing that agents augment their SR learning with a replay buffer that can replay experienced transitions to update the SR offline (Gardner et al.,2018; Momennejad, Russek et al., 2017). In fact, these two explanations might be closely related: In the neural network implementation of the Kalman filter introduced by Dayan and Kakade (2001) and applied to Kalman TD by Gershman (2017), the covariance matrix is approximated by a recurrent layer. Given a feature vector, the network activates other features whose weights positively covary with the weights of the currently activated features and it deactivates features whose weights negatively covary. This process can be seen as a covariance-based memory retrieval process similar to an attractor network. The “replay” process in this model amounts to a covariance-based memory retrieval process, similar to an attractor network: given the current feature vector, features whose weights positively covary with the weights of the active features are activated, and features whose weights negatively covary are deactivated. Applying a prediction error update to this ‘replay vector’ approximates the Kalman TD algorithm (Gershman, 2017). This is different from experience replay, in which experienced sequences of states are replayed (Lin, 1992), and simulated experience, in which possible experience sequences are generated from a transition model (Momennejad, Otto et al., 2017; Sutton, 1991); rather, multiple states are potentially activated and updated simultaneously (like the form of reactivation described in (Manning, 2021)).

Thus, the Kalman filter model suggests a biologically plausible implementation of a rapid covariance-based replay mechanism that would capture these results. It is interesting to note that Momennejad, Russek et al. (2017) also found that transition revaluation was associated with longer reaction times than reward revaluation. Under our interpretation, this could be attributable to either uncertainty leading to longer reaction times or the recurrent dynamics of settling into an attractor state under the biological implementation. Note that another related model achieves replanning after transition changes without replay by adding a low-rank correction matrix to the original representation (Piray & Daw, 2021b).

A further reduction in uncertainty about the SR could be achieved using offline inference or smoothing. The Kalman filter’s uncertainty estimates could be an interesting measure for determining which states should be replayed (Evans & Burgess, 2019). An alternative metric for the utility of replaying a specific state, suggested by Mattar and Daw (2017), is the product of a gain and need term, where the need term corresponds to the SR and the gain term quantifies the net increase in value expected after a policy change in a given state. This latter measure does not explicitly take into account uncertainty, but such a term might be approximated using the *value of information*, which can be computed from uncertainty estimates (Dearden et al., 1998). In addition to prioritizing sequences of replayed states, information about uncertainty may also be used for exploration of states with high uncertainty (see Malekzadeh et al., 2022, for an application of Kalman SR in active learning). Replay and exploration both depend on an ability to generate sequential samples and correspond to different optimal sampling regimes, possibly mediated by entorhinal grid cells (McNamee et al., 2021).

For the context model, another interesting avenue for further research is to investigate whether we can understand replay as offline inference (i.e. smoothing) in the case of multiple maps. In switching Kalman filters, smoothing does not only sharpen the posterior of the within-mode continuous latent variable, it also makes the posterior over modes more precise (Barber,2012). In this context, replay could serve the function of better separating out different maps from each other, or alternatively, to merge maps where this is appropriate. Guo et al. (2020) found evidence that a single coherent map is being built during sleep. In Lever et al. (2002), maps for environments of different geometries differentiate within trials but they get more similar again between trials.

### 4.2 Effect of shock on context inference

In the proposed model, inferred context changes are induced by large magnitude state prediction errors. This context allocation mechanism allowed the agent to separate associations made in different environments, or indeed in the same environment with different future behaviour, such as in the fear conditioning experiments of Winocur et al. (2009). However, if context changes are induced by large prediction errors, one would expect the negative reward prediction error associated to the shock in fear conditioning itself to be a driver of context switching. This raises the question how this type of prediction error would affect the model presented here.

Consistent with the idea that negative reward prediction error should be a driver of context change, it has been observed that electrical shocks in fear conditioning experiments can induce hippocampal place cell remapping (Moita et al., 2004). Intriguingly, however, this remapping was much stronger when animals learned that the environment itself was predictive of the negative reinforcement (context condition). By contrast, when the shock was paired with a specific auditory cue (cue condition, as was also the case in Winocur et al., 2009), only a small subset of the place cells showed remapping. As noted by the authors, one explanation for that disparity could be that the cells that remap are more involved in acquiring the new contextual associations than non-remapping cells. If this were the case, the cue condition would result in a re-coding only of the cue itself because the cue is what requires a rapid new association. The small amount of remapping that was still observed in this condition can be explained by the animals still acquiring some association between the environment and the shock. In the context condition, the new association is made with the place itself, requiring the local place cells to remap.

What would this look like in the model? When the negative reinforcement is applied, the prediction error drives a switch of context. In this new contextual representation, the features that are active at that moment acquire the association with the shock. In the cue condition, this association will mostly be with the cue, because this most salient cue will overshadow associations with active place cells. In the context condition, when no cue is present, this association will be with the place cells directly. On subsequent arrivals to the environment, the agent will have to infer which context applies. In the cue condition, the fearful context will have lower likelihood until the cue appears. In the context condition, any location that predicts the location where the shock arrived is consistent with the fearful context, driving a higher likelihood for the fearful condition (and thus remapping) for the whole environment. This interpretation means that if there are non-cued shocks that happen in very specific locations of a maze environment (with only low predicted occupancy from other parts of the environment), remapping should mainly be observed in that part of the environment. Interestingly, Schuette et al. (2020) showed that hippocampal neurons encode space at a finer scale following fear memory acquisition and that this effect is strongest near the shock grid. In our model, regions of the environment where the predicted occupancy of the shocked state is low should not show remapping.

### 4.3 Limitations and alternative models

We make several assumptions in order to make this model tractable. Firstly, the observation noise is assumed to be white (i.e. independent per time step). Since this assumption does not hold in many cases, we have included an analysis of the white noise assumption and an alternative model in Appendix B. Secondly, following the value estimation method described by (Geist & Pietquin, 2010a), we chose a random-walk model for describing the evolution process on the SR parameters. With this identity evolution model, all inference burden is put on the observation process. This means that Kalman TD is simply a reinterpretation of TD learning, i.e. a model-free way to estimate the SR. Given this evolution model, and assuming independent noise, we could make the assumption that the parameters for each successor feature (i.e. each column of the weight matrix) were independent such that, effectively, the Kalman SR model consists of *N* independent filters. Furthermore, since the evolution of the covariance matrix is independent of the prediction errors, the covariance matrix corresponding to each column was the same. Of course, in reality there do exist dependencies between the different columns of *M*. For example, in the tabular case, visiting any particular state more than expected means that all other states will be visited less than expected. A more sophisticated evolution model could exploit these dependencies; however, this would break the independence assumptions and thereby increase the computational burden. Inference in the switching Kalman filter is generally intractable, and we have chosen for a particle filter based approximation in our simulations. The experiments we modelled here do not speak to one or another form of approximate inference, but this is an interesting avenue for further research. Interestingly, trial-by-trial fluctuations and sudden changes are well-described by particle filter algorithms with a small number of particles, suggesting that the brain may indeed use sequential Monte-Carlo sampling (Daw & Courville, 2007).

A different way to combine the SR with uncertainty was proposed by Janz et al. (2018). In their approach, the SR *ψ* is approximated using temporal difference methods, without taking into account uncertainty, but the reward weight vector **w** is estimated using Bayesian linear regression. The authors use these reward uncertainty estimates to balance exploitation and exploration and show that their method is effective on a set of exploration benchmarks. Since the uncertainty estimates are not used to alter the updates of the SR itself, this model would not display behaviours modelled here such as transition revaluation. To allow more naturally for non-stationary reward functions, it would be interesting to swap the Bayesian linear regression for a Kalman filter. In other related work, (Madarasz, 2019) use a hierarchical model over multiple context-dependent SRs, using a sticky CRP prior to explain aspects of hippocampal activity in spatial environments. This work does not include a representation of within-context uncertainty, as captured by the Kalman filter in our model.

### 4.4 Suggested neural information processing architecture

While we have explicitly focused on modelling behaviour in this paper, a complete account will of course need to incorporate the brain regions involved in different parts of the model. We hypothesise that the different SR maps are encoded by the hippocampus, which shows many resemblances to the SR (Stachenfeld et al., 2017). For example, the firing fields of hippocampal place cells show experience-dependent skewing, consistent with a prediction of future locations (Mehta et al., 2000). Neuroimaging studies have furthermore shown predictive coding of non-spatial states (Garvert et al., 2017; Schapiro et al., 2016), although a recent direct test of the SR proved inconclusive about the role of the hippocampus (Russek et al., 2021). Accordingly, we propose that the prediction error-mediated switching between contextual SR maps corresponds to hippocampal remapping (Sanders et al., 2020). In addition, the OFC has previously been shown to be involved in predicting both reward outcomes (Gottfried et al., 2003; Schoenbaum et al., 1998) and sensory events (Chaumon et al., 2014) and is crucial for learning the stimulus-stimulus associations in Hart et al. (2020) (Figure 5). Accordingly, Wilson et al. (2014) have proposed that the OFC encodes a cognitive map of task space.

As for the prediction error used to update the SR, evidence from optogenetic studies in rodents suggests that these could be encoded by a population of dopamine neurons in the ventral tegmental area (VTA, see Figure A1; Gardner et al., 2018; Sharpe et al., 2017). Dopamine neurons are furthermore known to modulate the hippocampus, which in turn projects to the striatum (Lisman & Grace, 2005). Taken together, this suggests an information processing architecture in which SR maps are encoded in the hippocampus and OFC and updated by dopaminergic modulation. In this hypothesis, the striatum could compute values from the SR, feeding into action selection.

### 4.5 Conclusions

In conclusion, we introduced a model of reward prediction with the SR in which a distribution over SR weights is estimated using Kalman TD. In the model, the appropriate context is chosen based on how well a certain set of SR parameters serves for predicting the current observations. This model captures several learning phenomena, including the effects of context pre-exposure on learning and generalisation, the effects of reward devaluation after preconditioning and the context-specificity of memories. This paper demonstrates that these hitherto unconnected themes in animal learning can be unified under a single model.

## Acknowledgements

We thank Athena Akrami, Peter Dayan and Steven Hansen for helpful comments. JPG received a PhD studentship from the Gatsby Charitable Foundation and the Wellcome Trust. SJG received funding from Air Force Office of Scientific Research (FA9550-20-1-0413). NB was supported by a Wellcome Principal Research Fellowship, ERC Advanced Grant NEUROMEM, Wellcome Collaborative Award “Organising knowledge for flexible behaviour in the prefrontal hippocampal circuitry”.

## Appendix

## A Dopamine-dependent devaluation

Similarly to the results of Hart et al. (2020) discussed in the main text (Figure 5), our model captures the findings of Sharpe et al. (2017), which show that the sensitivity to reward devaluation after preconditioning is dependent on dopamine transients. To show this, the authors designed an experiment similar to that of Hart et al. (2020). It started with a preconditioning phase, during which associations were learned between pairs of neutral stimuli, followed by a conditioning phase during which a neutral stimulus was paired with a food reward (Figure A1A). After this conditioning phase, the food reward was devalued in one group but not another. To show that learning the associations between neutral stimuli was mediated by dopamine, there were two preconditioning phases, during which the authors applied a “blocking” (Kamin, 1967) design. They established that the preconditioned associations where only learned when learning was unblocked by optogenetically stimulating dopamine neurons during learning. A key finding of this experiment was that animals’ responding to the unblocked preconditioned cues (C) was sensitive to the subsequent devaluation of the food reward (Figure A1B, see also Hart et al., 2020). Thus, reward devaluation can alter stimulus-stimulus associations that were learned through the activation of dopamine neurons.

Gardner et al. (2018) simulated this experiment using an SR model and found that, while the SR accommodates many results found by Sharpe et al. (2017), in this particular experiment a standard SR agent is not sensitive to the reward devaluation (Figure A1B). As in Figure 5, this is because in the TD SR, only stimuli that directly predict reward will change value after devaluation, and C was never directly associated with food. Gardner et al. (2018) therefore simulated the task with an SR model endowed with the ability to simulate offline experience, which allowed the model to be sensitive to devaluation. As with the Hart et al. (2020) experiment, Kalman SR is sensitive to this devaluation paradigm without the need for adding an offline replay mechanism.

**Figure A1:**
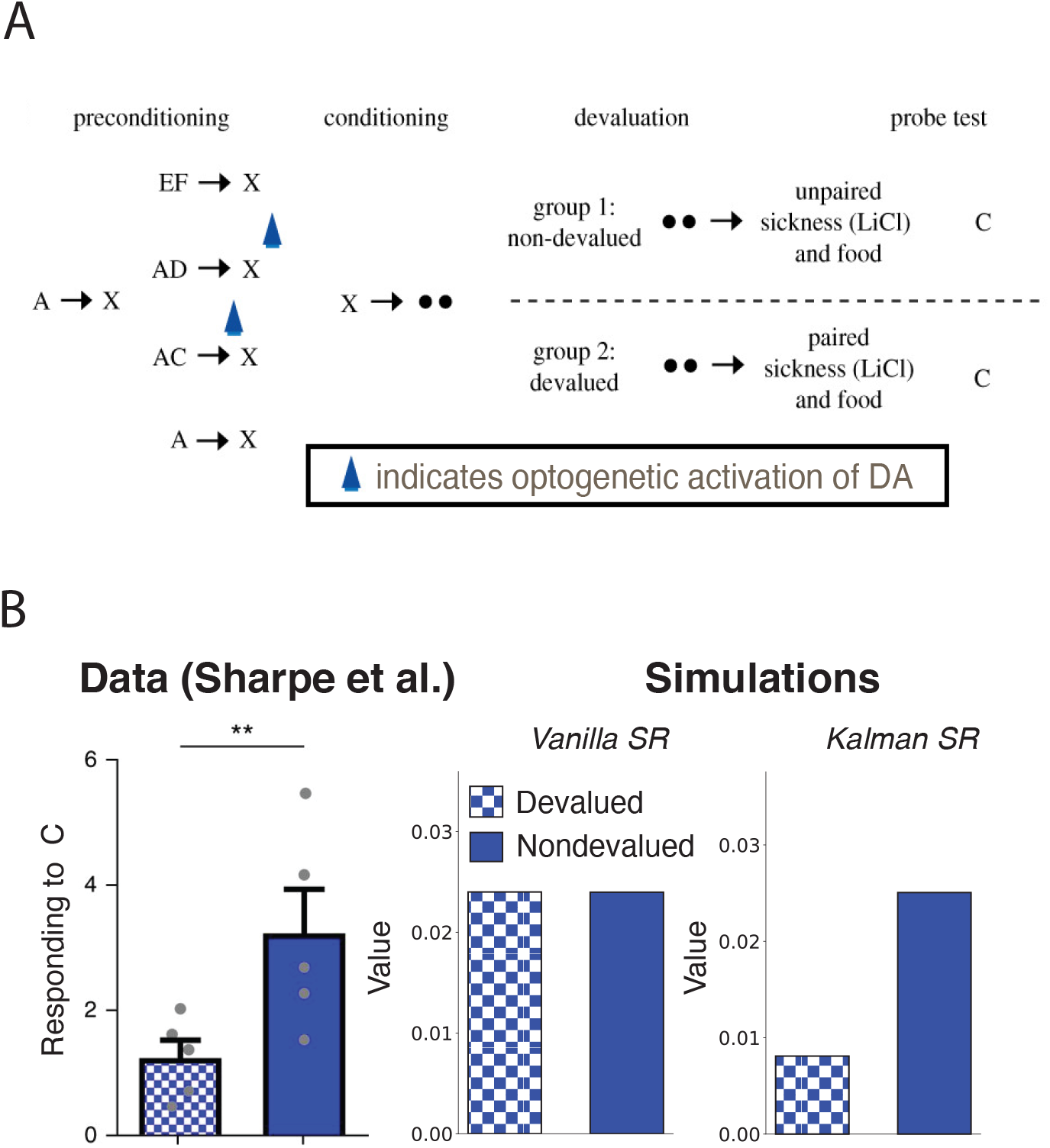
Devaluation experiment by Sharpe et al. (2017). **(A)** Experimental design. During the initial preconditioning phase, one neutral stimulus always precedes a second (*A* → *X*), after which the same preceding stimulus is compounded with a second predictor stimulus to precede the predicted stimulus (e.g. *AC* → *X*). The initial *A* → *X* pairing blocks learning about the second association, but this blocking is prevented by optogenetic activation of dopamine neurons during learning (Sharpe et al., 2017). After the conditioning phase, the food is paired with lithium chloride (LiCl) to induce sickness. Letters denote different neutral stimuli, black circles indicate food reward. **(B)** Data and simulation results show that, like animals and unlike TD-SR, Kalman SR shows sensitivity to devaluation in this paradigm.

## B The white noise assumption of Kalman TD

The basic version of the Kalman TD algorithm introduced in the main paper was derived based on the simplifying assumption that the observation noise is white (independent per time step). In reality, this is only the case when the transitions are deterministic. In that deterministic case, optimal update of the weights can be derived, resulting in the Kalman TD algorithm used in the paper (Geist & Pietquin, 2010b). In most cases, however, the successive uncertainty terms cannot be treated as independent because they are related by the way in which the agent moves through the world. When the transitions are stochastic, the expectation over successor states given the current state (and action, if applicable) must be considered. When this is not done, and the original Kalman TD cost function is applied to tracking a state value function *V*, it can be analytically shown that this leads to the following bias (Geist & Pietquin, 2010b):

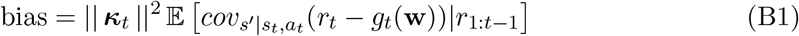

with 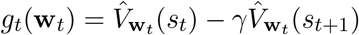, and ***κ*** denoting the Kalman gain.

This bias is inherent in applying Kalman TD in a stochastic setting to making predictions about any kind of cumulant, of which a reward function is only one example. Therefore, the same issue arises when estimating the SR, but for simplicity we discuss the value tracking case here. We discuss the issue of bias as well as a solution with an alternative noise model briefly here. For a more extensive discussion including derivations of the bias and the alternative noise model we refer the reader to Geerts (2021) and Geist and Pietquin (2010b).

To alleviate the issue of bias in Kalman TD, Geist and Pietquin (2010b) introduced a coloured noise model that was first introduced by (Engel et al., 2005) in Gaussian Process TD. The key idea is to replace the white observation noise in the generative model by a “coloured” observation noise, i.e. a noise that is not independent per time step. As shown by Geist and Pietquin (2010b), this involves extending the parameter vector **w** to include the observation noise, such that the observation noise will be estimated from data in the inference process. The computational complexity of the resulting algorithm, extended Kalman TD (XKTD), is the same as for the original Kalman TD because the parameter vector is extended with two scalars. However, this coloured noise estimation induces some memory effects which means that XKTD cannot be applied to off-policy evaluation.

In order to empirically asses how damaging the white noise assumption is, we now compare KTD’s value estimates to the true (unbiased) value, approximated by Monte Carlo (MC) sampling. In MC sampling, value is estimated by simply averaging sample returns across episodes (Sutton & Barto, 1998). For completeness, we also compare these to XKTD estimates.

We evaluated both algorithms on a simple chain MDP (adapted from Brockman et al., 2016). The MDP has seven non-absorbing states, arranged linearly from state 0 to 6. Making a right move in the final state leads to an absorbing state. The agent can move to the left or right and receives a reward of −0.2 for every step except in the absorbing state, where it receives a reward of 10. The stochasticity in the state transitions will come from the policy, which can be defined by a single parameter *P*(*R*), for the probability of making a step to the right (*P*(*L*) = 1 – *P*(*R*)).

**Figure B1:**
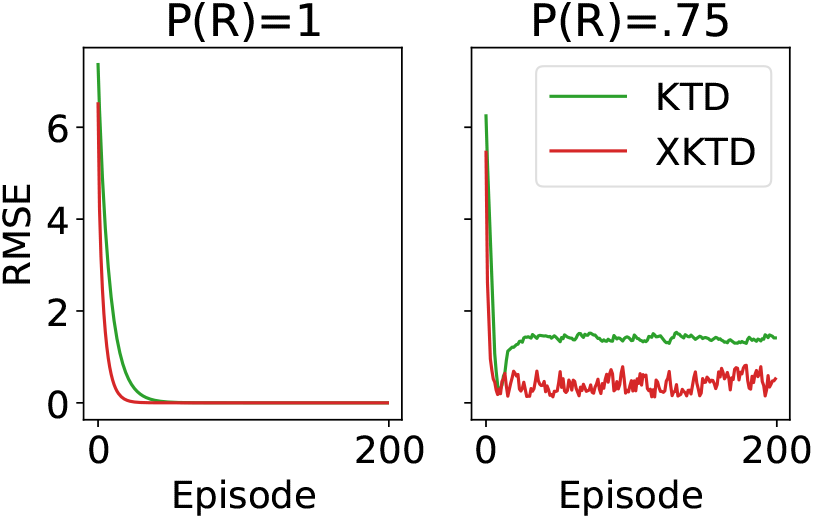
Root mean square error after each episode for an example run of KTD and XKTD in a deterministic (left) and stochastic (right) MDP.

We ran KTD and XKTD on this domain for 200 episodes, with a deterministic optimal policy (*P*(*R*) = 1) and with a stochastic policy (*P*(*R*) = 0.75), computing after each episode the root mean square error (RMSE) between the true value function (as estimated by Monte Carlo) and the algorithm’s value estimate (Figure B1). With deterministic transitions, both algorithms converge to the same low error (left panel), but with stochastic transitions, KTD converges to a wrong value, maintaining higher error, consistent with the analytically derived bias (equation B1). Figure B2 shows the posterior distribution over value after the example run of 200 episodes for both KTD and XKTD, overlaid with the actual, sampled returns. Indeed, the mean of the posterior for KTD is consistently off, while the XKTD posterior is closer to the true mean.

To quantify how damaging the deviations are as a function of stochasticity of the environment, we then varied *P*(*R*) from 1 to 0.5 (completely random transitions), running KTD and XKTD for 200 episodes, repeated this 10 times for each value of *P*(*R*) and computed the RMSE, which is shown in Figure B3. As could be seen theoretically, the bias grows as the environment is more stochastic. In addition, the bias is significantly reduced for XKTD, although even for the latter algorithm the bias grows with higher stochasticity.

**Figure B2:**
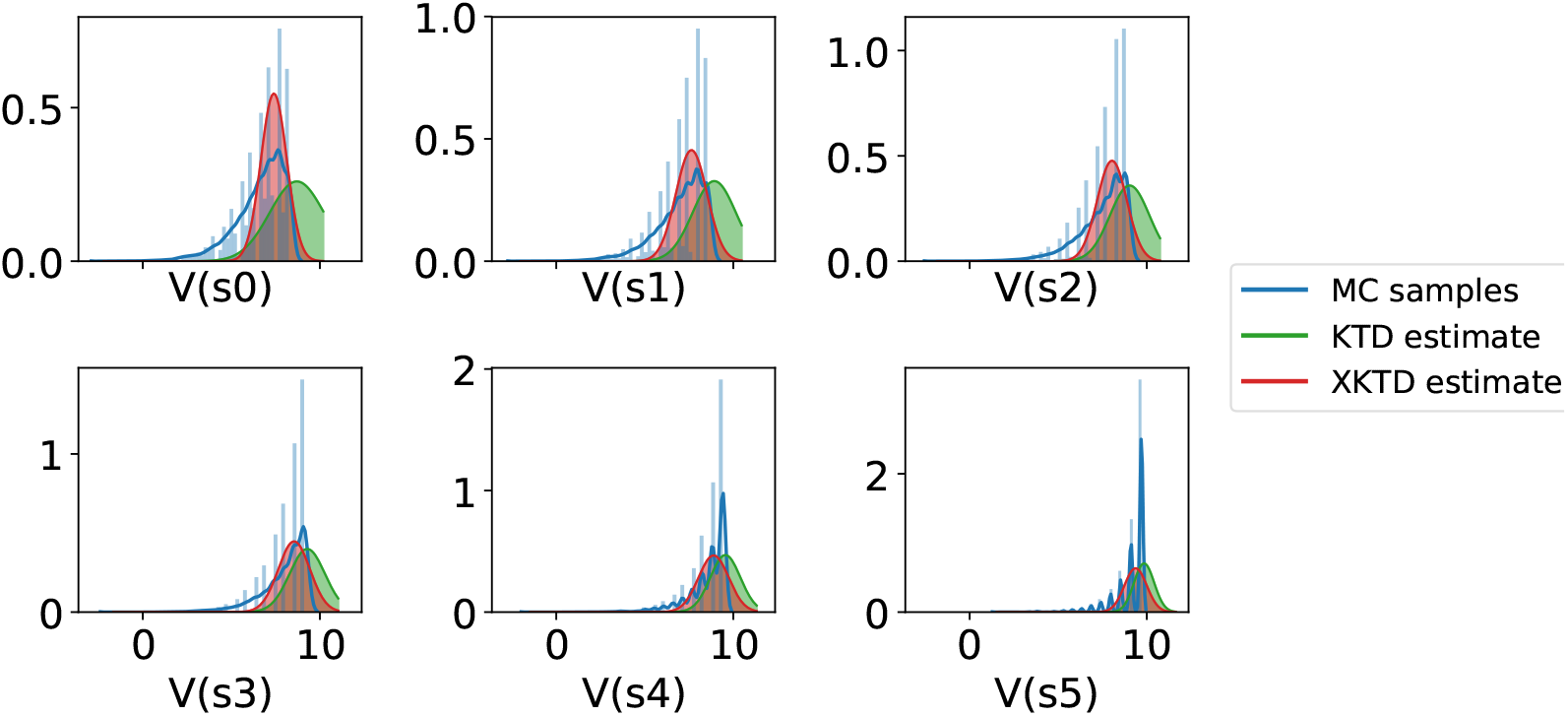
Posterior distributions over value after an example run of 200 episodes for KTD and XKTD, overlaid with the true, sampled distribution of returns.

**Figure B3:**
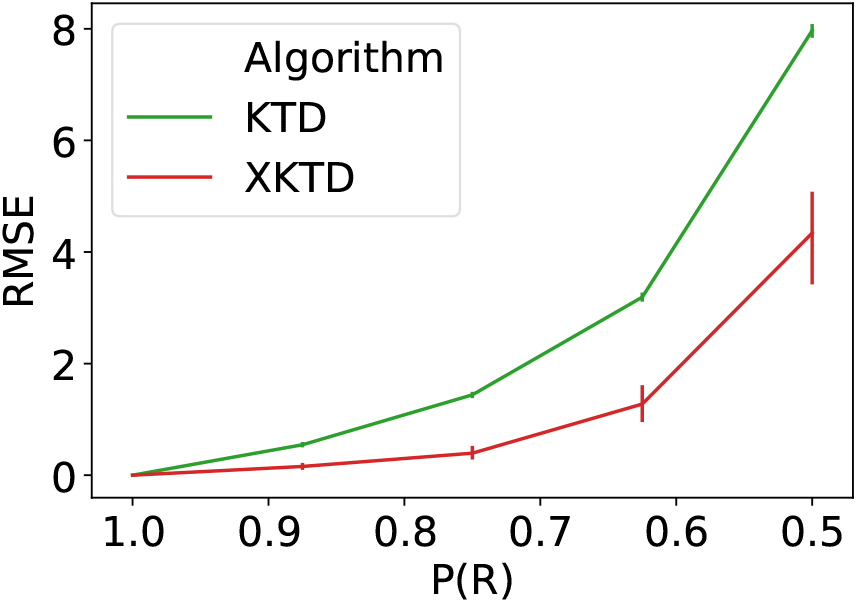
Comparing KTD and XTKD on the linear track environment. RMSE after 200 episodes is plotted as a function of stochasticity of the environment (*P*(*R*) = 0.5 corresponds to maximum entropy / randomness). Error bars show 95% confidence intervals.

Thus, the Kalman TD algorithm incorrectly treats successive observation noise terms as “white” (independent from each other), while they are related because of the way the agent moves through the world. Any stochasticity in the transitions will therefore bring about a bias in the KTD estimates. This problem can be alleviated using XKTD, in which the hidden parameter vector is extended such that the observation noise, which is now assumed to be coloured, can be estimated online. However, as shown in Figure B3, even XKTD leads to biased estimates under high stochasticity. This is because, while the assumptions are less strong than for KTD, XKTD still incorrectly assumes that the successive residuals are independent from each other.

In conclusion, KTD’s uncertainty estimation is incorrect for many realistic MDPs, but this can be remedied by extending KTD with coloured noise estimation, without adding significant computational or memory complexity.

